# PepRePs: Peptide-Retargeted Phosphatases Designed with Generative Language Models

**DOI:** 10.64898/2026.10.05.756719

**Authors:** Lauren Hong, Tai-Chen Ho, Yi-Shiuan Tseng, Sophia Vincoff, Tong Chen, Thomas Connor Donahue, Matthew P. DeLisa, Po-Han Chen, Pranam Chatterjee

## Abstract

Phosphorylation is a fast, reversible layer of post-translational control that directly governs protein activity, signaling flux, and cell state. Kinases and phosphatases collectively establish these phosphorylation states, but their dysregulation drives cancer, neurodegenerative diseases, and developmental disorders. Selectively manipulating the phosphorylation of individual proteins in cells remains challenging, particularly for disordered substrates. Here, we introduce **Pep**tide-**Re**targeted **P**hosphatase**s** (**PepRePs**), a genetically encoded system that fuses a peptide binder to a serine/threonine phosphatase subunit to localize phosphatase activity to a target protein. We first establish the PepReP architecture using characterized tau-binding peptides, then apply protein language model-guided peptide design to target Rab8A, PTEN, and STAT3. Across endogenous and exogenous targets, PepRePs reduce phosphorylation, with downstream effects on tau aggregation, Rab8A-associated protein interactions, and cellular proliferation following PTEN and STAT3 targeting. Sequence-based peptide design enables targeting without structural inputs, supporting application to substrates with disordered regions that challenge conventional structure-based binder design. By coupling programmable peptide binders with modular phosphatase subunits, PepRePs enable targeted control of protein phosphorylation to probe cellular signaling and develop strategies for correcting disease-associated protein dysfunction.

## 1 Introduction

Cell state depends not only on gene expression but also on rapid, reversible post-translational control of protein activity, localization, and interaction. The addition or removal of phosphate on serine, threonine, and tyrosine residues alters protein conformation, charge distribution, and binding surfaces, propagating changes in signaling output and cell behavior [1–4]. Dysregulated phosphorylation is causally linked to cancer, where constitutively active oncogenic kinase signaling drives proliferation and survival [5]; to neurodegenerative disorders such as Alzheimer’s disease, where hyperphosphorylated tau loses its capacity to stabilize microtubules [6]; and to Huntington’s disease, where the phosphorylation state of huntingtin modulates its aggregation and toxicity [7]. Current strategies to manipulate phosphorylation primarily target the enzymes that establish these states. Kinase and phosphatase inhibitors and activators consequently perturb multiple substrates across a signaling network [8, 9]. More targeted approaches recruit phosphatase activity to individual substrates through bifunctional small molecules such as phosphorylation-targeting chimeras (PhosTACs) [10, 11], short peptide motifs such as dephosphorylation targeting chimeras (DEPTACs) [12–14], or genetically encoded nanobody-phosphatase fusions such as AdPhosphatase [15]. These approaches provide substrate-level control, but their targeting architectures remain constrained by the availability of suitable small-molecule binding sites, linear motifs, or folded epitopes. This limitation is particularly relevant for phosphosites embedded within intrinsically disordered regions (IDRs), where the local sequence may not form a stable structure suitable for conventional binder design [16].

Peptide binders provide a compact targeting modality that can engage extended protein surfaces while remaining amenable to genetic fusion with enzymatic domains [16]. Genetically encodable proteome-editing systems have established that short peptide binders can serve as modular intra-cellular targeting elements: peptide-guided ubiquitination systems (ubiquibodies, or uAbs) recruit E3 ligases to induce selective protein degradation [17–20], while peptide-guided deubiquitinases (deubiquibodies, or duAbs) enable targeted protein stabilization [21]. These systems establish that peptide-guided recruitment can redirect enzymatic activity toward selected intracellular substrates. Extending this approach to phosphorylation requires peptide binders that can be generated across diverse protein sequences, including targets that lack defined structures for structure-based methods such as RFdiffusion, BindCraft, or BoltzGen [22–24]. Protein language models (pLMs) provide a sequence-based route to this design problem by learning protein representations without explicit structural inputs [25, 26]. SaLT&PepPr, PepPrCLIP, and PepMLM have demonstrated sequence-based generation and prioritization of short peptide binders [18–20], including binders used to construct functional uAbs and duAbs [21]. More recent methods such as moPPIt and SOAPIA further enable motif-level and isoform-aware peptide targeting [27, 28]. These advances provide a foundation for programming peptide-guided phosphatase recruitment across structurally diverse targets.

Phosphorylation regulates protein function and signaling state on timescales that complement transcriptional and protein-abundance control [3, 29]. Here, we introduce **Pep**tide-**Re**targeted **P**hosphatase**s** (**PepRePs**), a genetically encodable system that fuses pLM-designed peptide binders to serine/threonine phosphatase subunits to enable programmable, selective dephosphorylation of endogenous proteins. We show that PepRePs reduce site-specific phosphorylation of five disease-relevant targets, tau [30], Rab8A [31], STAT3 [32], and PTEN [33], and produce downstream effects on cellular phenotypes associated with these phosphorylation events. By coupling sequence-based peptide design with modular phosphatase recruitment, PepRePs provide a genetically encoded approach for directly perturbing phosphorylation states within intracellular signaling networks.

While peptide-guided ubiquitination and deubiquitination systems (uAbs and duAbs) regulate protein abundance, phosphorylation governs protein function on faster timescales and often dictates signaling state without altering expression levels [3, 29]. Here, we introduce **Pep**tide-**Re**targeted **P**hosphatases (**PepRePs**), a genetically encodable platform that fuses pLM-designed peptide binders to serine/threonine phosphatase subunits, enabling programmable, selective dephosphorylation of endogenous proteins. We show that PepRePs dephosphorylate disease-relevant targets, including tau [30], Rab8A [31], STAT3 [32], and PTEN [33], reducing site-specific phosphorylation and producing downstream functional effects on cellular behavior.

## 2 Results

### 2.1 Established peptides guide the engineering of PepRePs for targeted tay de-phosphorylation

Previous work has established that short peptide binders can be fused to enzymatic domains to enable targeted protein degradation [17] or stabilization [21]. Building on this paradigm, we designed PepRePs by fusing target-binding peptides with catalytic serine/threonine phosphatase subunits for recruiting phosphatase activity to a selected protein (Figure 1A; Supplementary Table S1). We first evaluated this approach using tau, a microtubule-associated protein that becomes hyperphosphorylated and prone to aggregation in Alzheimer’s disease (Figure 1B) [12, 34]. We selected three previously charcterized tau-binding peptides as targeting elements. TB1 was previously developed for tau dephosphorylation in Alzheimer’s disease mouse models [35], whereas TB2 and TB3 were reported to recognize tau fibrils and reduce phosphorylated tau levels in cellular and *in vivo* models (Supplementary Table S2) [12]. These binders provided a set of independently characterized binders with which to evaluate phosphatase recruitment to tau.

**Figure 1:**
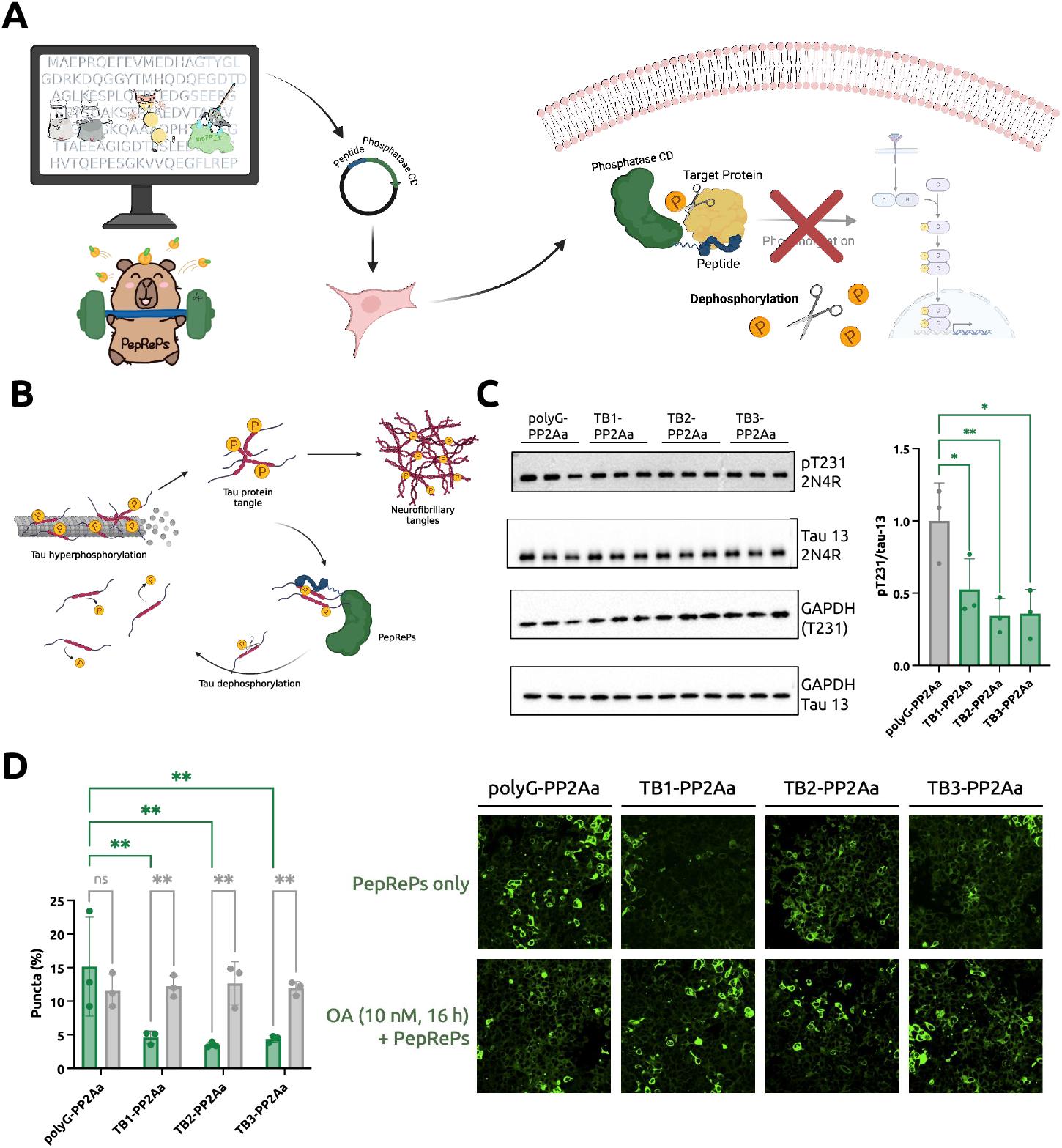
Tau-targeting PepRePs reduce tau phosphorylation and aggregation-associated puncta. A) Genetically-encodable architecture of peptide-recruiting phosphatases. B) Schematic of tau targeting the relationship between tau phosphorylation and aggregation. C) Tau pT231 dephosphorylation by PepRePs (using PP2Aa subunit) in tau-expressing HEK293 Flp-In Trex cells. D) Confocal microscopy of tau-associated puncta showing PepRePs expression and treatment with the phosphatase inhibitor okadaic acid (OA). Statistical analysis was performed using an ordinary one-way ANOVA with Dunnett’s multiple comparisons test for densitometry analysis and a two-way ANOVA with Šídák’s multiple comparisons test for tau-associated puncta analysis. All analyses were performed using GraphPad Prism 11 software, with calculated *p* values represented as follows: ^*^, p *≤* 0.05; ^**^, p *≤* 0.01. Please refer to source data for numeric p values.

We next evaluated whether the choice of phosphatase subunits influenced PepRePs activity toward tau dephosphorylation. We tested the catalytic subunit of PP1 (PP1c), the regulatory and catalytic subunits of PP2A (PP2Aa and PP2Ac, respectively, and the catalytic subunits of PP4, PP5, and PP6 (PP4c, PP5c, and PP6c). These phosphatases represent distinct phosphoprotein phosphatase (PPP) architectures in which substrate recognition and enzyme activity are regulated through associated targeting subunits or regulatory domains [36–40]. Among the phosphatase configurations tested, the PP2Aa-containing PepRePs produced the most pronounced reduction in tau pT231 phosphorylation (Figure 1C) [41]. This result is consistent with the established role of PP2A as a major tau-directed phosphatase in the brain [34, 42, 43], and identified PP2Aa as an effective catalytic component for subsequent tau-targeting experiments. The reduction in tau phosphorylation was accompanied by a decrease in tau-associated puncta in a cellular tau seeding assay, indicating that PepReP-mediated dephosphorylation altered a downstream phenotype associated with tau aggregation (Figure 1D) [44].

To determine whether this phenotype depended on phosphatase activity, we treated cells expressing the tau-targeted PepRePs with okadaic acid (OA), a phosphatase inhibitor. OA treatment restored tau phosphorylation and increased tau-associated puncta, supporting a phosphatase-dependent mechanism for the observed reduction in tau aggregation (Figure 1D). Together, these results establish that peptide-guided recruitment of a serine/threonine phosphatase can reduce phosphorylation of an intracellular tau substrate and produce a corresponding cellular phenotype. The ability to evaluate different targeting peptides and phosphatase subunits within the same architecture provides a modular basis for adapting PepRePs to additional phosphoprotein targets.

### 2.2 pLM-designed peptides enable targeted Rab8A dephosphorylation

To assess the programmability of PepRePs beyond tau, we next targeted Rab8A, a membrane-trafficking GTPase whose phosphorylation is regulated by the Parkinson’s disease-associated kinase LRRK2 [31, 45–47]. LRRK2 phosphorylates Rab8A at Thr72 within the switch II region, and this modification alters interactions with Rab8A effectors involved in membrane trafficking [31]. Recent studies further support Rab8A phosphorylation as a downstream consequence of pathogenic LRRK2 signaling in cellular models [46]. We selected candidate Rab8A-binding peptides using SaLT&PepPr by leveraging a known protein-protein interactions – specifically with partners Rab6-interacting protein 1 (residues 900-916, Rab8A_SnP_1) and Rabin8 (residues 189-208, Rab8A_SnP_2) – and identifying candidate interaction motifs from the sequences of Rab8A-interacting proteins (Supplementary Table S2) [19]. These peptides were fused to candidate phosphatase catalytic subunits to generate Rab8A-targeted PepRePs (Figure 2A; Supplementary Table S1).

**Figure 2:**
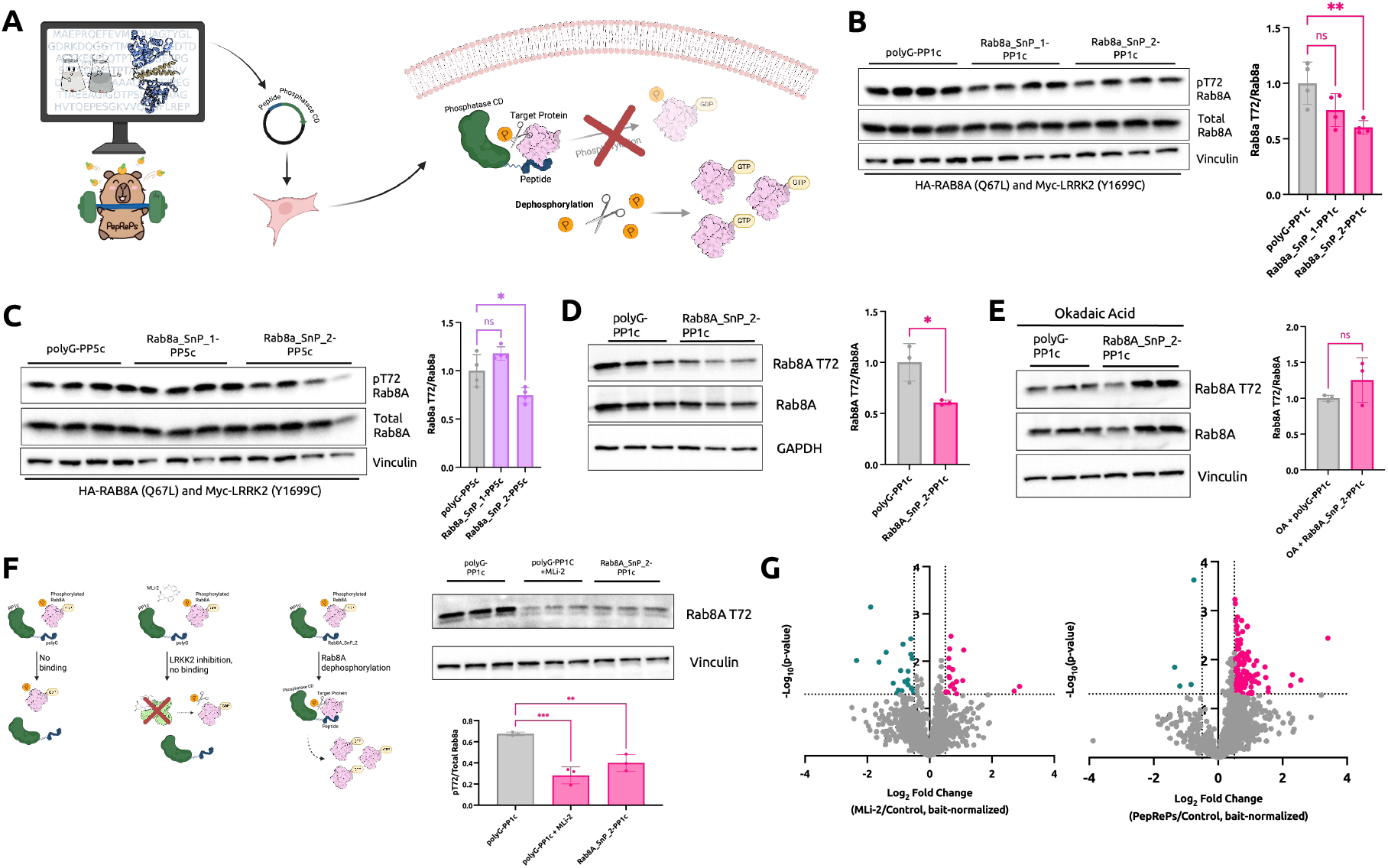
Rab8A-mediated PepRePs design. A) Genetically-encodable architecture for peptide-recruiting phosphatases. B) Rab8A-targeting PepRePs employing the PP1 catalytic subunit (PP1c). C) Rab8A-targeting PepRePs employing the PP5 catalytic subunit (PP5c). D) HA-tag pulldown of Rab8A showing differential expression of Rab8A Thr72 phosphorylation due to PepRePs treatment. E) HA-tag pulldown of Rab8A showing non-differential expression of Rab8A Thr72 phophorylation upon okadaic acid treatment. F) Comparison of targeted Rab8A dephosphorylation by the Rab8A_SnP_2-PP1c PepReP and pharmacological inhibition of LRRK2 with MLi-2. G) Volcano plots showing differentially regulated proteins following MLi-2 treatment compared to control (left) and Rab8A_SnP_2-PP1c PepRePs expression compared to control (right). Statistical analysis was performed using an ordinary one-way ANOVA with Dunnett’s multiple comparisons test and a two-tailed Student’s t-test. Analyses were performed using GraphPad Prism 11 software, with calculated *p* values represented as follows: ns, not significant, ^*^, p*≤* 0.05; ^**^, p *≤* 0.01. Please refer to source data for numeric p values.

We first screened the resulting constructs for their ability to reduce Rab8A Thr72 phosphorylation. Among the phosphatase configurations tested, PP1c-based PepRePs produced the most pronounced reduction in Rab8A pT72, as measured by phospho-specific immunoblotting, whereas the other catalytic subunits produced weaker effects (Figure 2B-C). We therefore selected the Rab8A_SnP_2-PP1c construct for subsequent characterization. To further validate targeted dephosphorylation of Rab8A in a protein-complex context, we performed HA-tag pulldown assays using HA-Rab8A to enrich Rab8A-associated complexes [44, 48]. Rab8A_SnP_2-PP1c reduced Rab8A pT72 in the enriched fraction, and this effect was abolished following treatment with okadaic acid (OA), supporting a requirement for phosphatase catalytic activity (Figure 2E). These results establish that peptide-guided recruitment of PP1c can reduce phosphorylation of Rab8A within an intracellular protein complex.

We next asked whether PepReP-mediated dephosphorylation of Rab8A could be achieved independently of pharmacological inhibition of its upstream kinase. LRRK2 is a principal kinase responsible for Rab8A Thr72 phosphorylation, and pharmacological inhibition with MLi-2 reduces Rab8A pT72 in cells [31]. Consistent with these studies, MLi-2 treatment reduced Rab8A Thr72 phosphorylation. Rab8A_SnP_2-PP1c produced a comparable reduction in Rab8A pT72 without directly inhibiting LRRK2, demonstrating that PepRePs can perturb the phosphorylation state of a kinase substrate through direct recruitment of a phosphatase (Figure 2F).

To determine whether these distinct perturbations produced similar effects on the Rab8A protein interactome, we performed quantitative proteomic analysis following Halo-trap enrichment of Rab8A-associated complexes. Although the proteomic responses to Rab8A_SnP_2-PP1c and MLi-2 were positively correlated (Pearson *r* = 0.67, Spearman *ρ* = 0.64; Supplementary Figure S1), the magnitude and composition of the responses differed substantially. MLi-2 treatment resulted in 16 significantly increased and 16 significantly decreased proteins, whereas Rab8A_SnP_2-PP1c resulted in 133 significantly increased and 3 significantly decreased proteins (Figure 2G). Thus, despite producing a similar reduction in Rab8A Thr72 phosphorylation, peptide-guided phosphatase recruitment and upstream kinase inhibition produced distinct changes in the Rab8A-associated proteome. These results demonstrate that PepRePs can directly alter the phosphorylation state of a defined kinase substrate while producing a proteomic response that differs from pharmacological inhibition of the upstream kinase.

### 2.3 *De novo* peptide design extends PepRePs to new targets

Finally, to demonstrate the programmability of our PepRePs platform, we *de novo* designed peptides with the target-conditioned algorithms PepMLM and moPPIt to PTEN and STAT3 [20, 27], two signaling proteins whose phosphorylation regulates distinct aspects of cellular function [32, 49]. PTEN Ser380 lies within the C-terminal phosphorylation cluster that regulates PTEN stability and catalytic activity [49]. Candidate PTEN-targeting peptides were fused to PP1c and screened for their ability to reduce PTEN pS380 relative to polyA-PP1c and untreated controls (Figure 3C). Multiple PTEN-targeting PepRePs significantly reduced PTEN pS380, with PTEN_moPPIt_2-PP1c producing a pronounced reduction relative to both controls. We next evaluated whether this reduction in PTEN phosphorylation was associated with a cellular phenotype. We expressed PTEN_moPPIt_2-PP1c and PTEN_PepMLM_3-PP1c in cells, and proliferation was measured over 48 hours. Both constructs reduced proliferation relative to untreated and polyA-PP1c controls, with PTEN_PepMLM_3-PP1c producing the larger reduction at later time points (Figure 3D).

**Figure 3:**
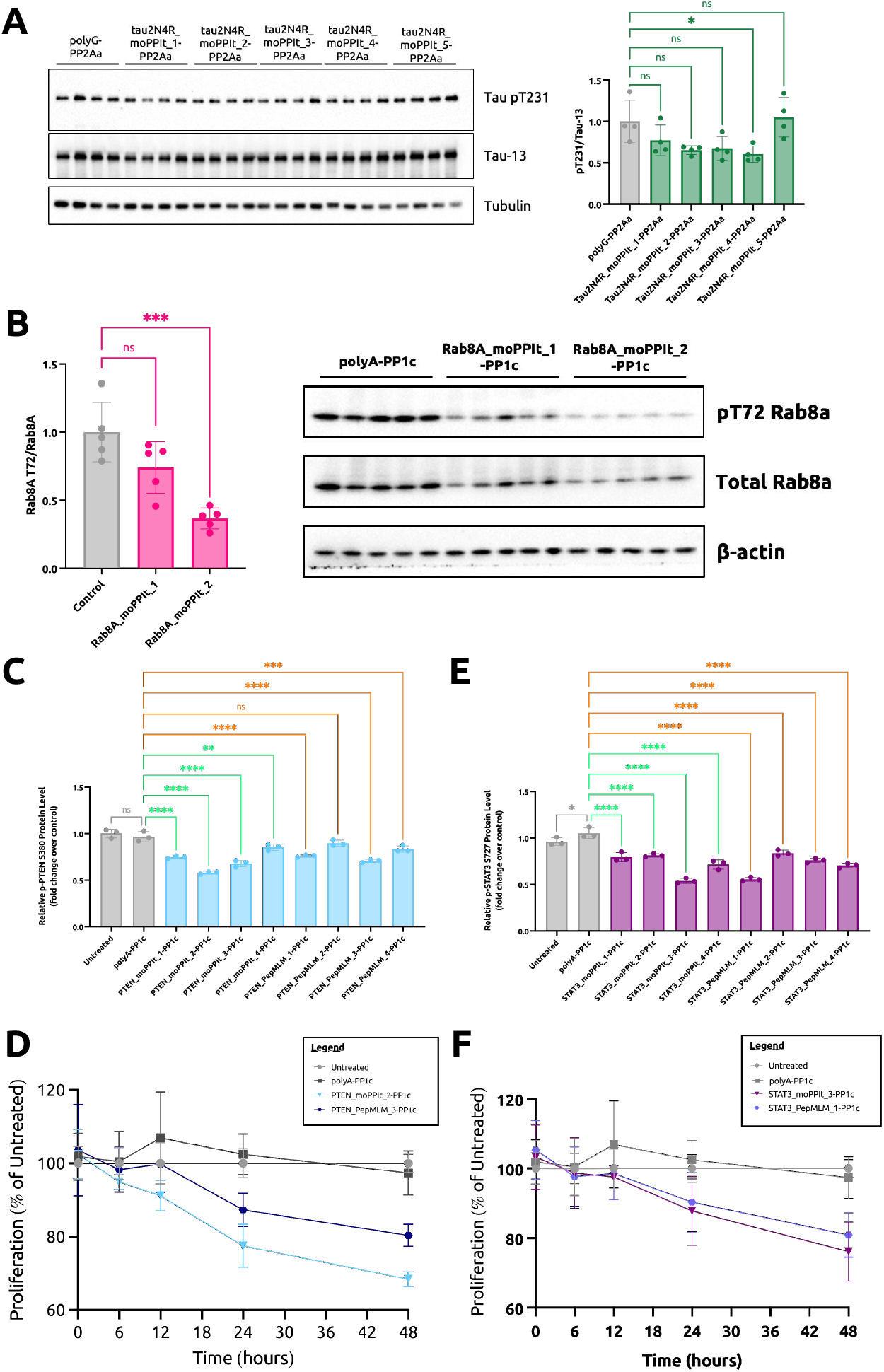
Extending PepRePs programmability. A) Tau-targeting PepRePs using moPPIt- and PepMLM-designed pepides. B) Rab8A-targeting PepRePs using moPPIt- and PepMLM-designed pepides. C) PTEN-targeting PepRePs using moPPIt- and PepMLM-designed pepides. D) MTS proliferation results of PTEN-designed PepRePs inducing decreased cell proliferation. E) STAT3- targeting PepRePs using moPPIt- and PepMLM-designed pepides. F) MTS proliferation results of STAT3-designed PepRePs inducing decreased cell proliferation. Statistical analysis was performed using an ordinary one-way ANOVA with Dunnett’s multiple comparisons test. Analyses were performed using GraphPad Prism 11 software, with calculated *p* values represented as follows: ns, not significant, ^*^, p *≤* 0.05; ^**^, p *≤* 0.01; ^***^, p*≤* 0.001; ^****^, p*≤* 0.0001. Please refer to source data for numeric p values.

We applied the same strategy to STAT3, a transcription factor whose Ser727 phosphorylation regulates transcriptional activity and cellular responses [32, 50]. Candidate STAT3-targeting peptides, designed by both PepMLM and moPPIt, were fused to PP1c and screened for their effects on pS727-STAT3 (Figure 3E). Multiple of these PepRePs significantly reduced pS727-STAT3 relative to polyA-PP1c and untreated controls, with STAT3_moPPIt_3-PP1c producing a pronounced reduction in phosphorylation. To determine whether targeted STAT3 dephosphorylation affected cellular behavior, we evaluated proliferation following expression of STAT3_moPPIt_3-PP1c and STAT3_PepMLM_1-PP1c. Both constructs reduced proliferation over 48 h relative to untreated and polyA-PP1c controls, with the effects becoming more pronounced at later time points (Figure 3F). Together, these experiments extend PepReP-mediated dephosphorylation to additional regulatory phosphorylation sites and demonstrate that targeted reduction of phosphorylation can produce measurable cellular phenotypes across distinct signaling proteins.

## 3 Discussion

In this work, we establish PepRePs as a simple, genetically-encoded strategy for directing serine/threonine phosphatase activity to defined intracellular proteins and reducing phosphorylation at disease-relevant sites. PepRePs reduced phosphorylation of four targets – tau, Rab8A, STAT3, and PTEN – across distinct cellular contexts and produced downstream changes in cellular phenotypes associated with the targeted phosphorylation events. The targeting elements spanned experimentally characterized peptides, peptides derived from known protein interactions, and *de novo* peptide binders generated from protein sequence. For STAT3 and PTEN, PepRePs enabled targeting without a pre-existing peptide binder or known binding partner as a starting point, demonstrating that peptide-guided phosphatase recruitment can be programmed from target sequence. This sequence-level targeting architecture complements existing approaches based on bifunctional small molecules, short linear motifs, or nanbody-phosphatase fusions [10, 12, 15] and provides a compact, genetically encodable mechanism for recruiting enzymatic activity to diverse protein targets [16]. By extending peptide-guided proteome editing from control of protein abundance to direct manipulation of phosphorylation-dependent protein fucntion, PepRePs provide a framework for perturbing intracellular signaling at the level of post-translational regulation.

The results also define the specificity boundary and design space of PepRePs. Peptide recruitment increases the local concentration of phosphatase activity near a target but does not establish aboslute phosphosite specificity, and our measurements do not determine whether additional sites on the target, associated proteins, or other phosphatase substrates are affected. This distinction is particularly relevant to the Rab8A experiments, where direct phosphatase recruitment and upstream LRRK2 inhibition produced overlapping but distinct changes in the Rab8A-associated proteome, while the interactome analysis did not resolve which changes arose directly from Rab8A dephosphorylation [51]. Systematic variation of peptide binders, phosphatase components, linker architectures, and expression levels, together with phosphoproteomic analysis, will further define the parameters governing activity and selectivity. Coupling this framework with language model-generated peptide binders [18–20] could enable scalable interrogation of previously unexplored phosphorylation sites and state-specific protein variants through binders designed for modified or mutant forms [27, 28, 52, 53]. More broadly, the PepReP architecture is not limited to phosphatases and can be extended to other PTM-modifying enzymes, including acetylases, glycosylases, and kinases, supporting a unified, programmable approach to post-translational regulation. Together, this work positions PepRePs as a general proteome editing platform that bridges generative modeling and experimental perturbation, enabling principled manipulation of post-translational state space with implications for both therapeutic development and foundational studies of cellular decision-making.

## 4 Methods

### Binder Design

The TB1, TB2, and TB3 peptides were derived experimentally as described in literature [12]. Novel binding peptides designed in this study were generated by the previously described SaLT&PepPr [19] (https://huggingface.co/ubiquitx/saltnpeppr), PepMLM [20] (https://huggingface.co/ChatterjeeLab/PepMLM-650M), and moP-PIt algorithms [27] (https://huggingface.co/ChatterjeeLab/moPPIt). All binder sequences can be found in Supplementary Table S2.

### Generation of Plasmids

All PepRePs plasmids were generated from the standard pcDNA3 vector, harboring a cytomegalovirus (CMV) promoter. An Esp3I restriction site was introduced immediately upstream of the phosphatase subunits (PP1c, PP2Aa, PP2Ac, PP4c, PP5c, and PP6c) and flexible GSGSG linker via Gibson Assembly using synthesized gene fragments (Azenta). For target-specific PepRePs assembly, oligos for candidate peptides were annealed and ligated via T4 DNA Ligase into the Esp3I-digested duAb backbone. Assembled constructs were transformed into 50 *µ*L NEB Turbo Competent *Escherichia coli* cells, and plated onto LB agar supplemented with the appropriate antibiotic for subsequent sequence verification of colonies and plasmid purification (Azenta).

### Cell Culture

The HEK293T, HEK293 Flp-In, and MDA-MB-231 cells were maintained in Dulbecco’s Modified Eagle’s Medium (DMEM) supplemented with 100 units/mL penicillin, 100 mg/mL streptomycin, and 10% fetal bovine serum (FBS). The A549 cells were maintained in a 1:1 mixture of Dulbecco’s Modified Eagle’s Medium (DMEM) and Ham’s F-12K (Kaighn’s modification), supplemented with 100 units/mL penicillin, 100 mg/mL streptomycin, and 10% fetal bovine serum (FBS). For endogenous target screening in cell lines, pcDNA-PepRePs (750 ng) plasmids were transfected into cells as triplicates (3 *·* 10^4^/well in an 8-well chamber, or 1 *·* 10^4^/well in an 96-well plate) with Lipofectamine 2000 in Opti-MEM. Cells were harvested after 48 hours for subsequent cell fractionation and immunoblotting.

### Tau Seeding Assay

Tau-aggregated seeds were prepared by incubating tau 0N4R P301S monomers in 20 *µ*M heparin, 2 mM DTT and 1X protease inhibitors in PBS for 72 hours at 37^*°*^C while shaking at 220 rpm [11]. Prior to seeding, HEK 293 Flp-In T-Rex cells expressing tau variants were plated for 24 hours prior to tetracycline (2 *µ*g/mL) treatment to induce Halo-tau variants expression. Aggregated seeds were mixed with tau-seeding reagent composed of Lipofectamine 2000, Opti-MEM, and tetracycline prior to cell treatment for 48 hours. HEK 293 Flip-IN cells were transfected with PepRePs with Lipofectamine 2000 in Opti-MEM for 72 hours. Inhibitor-specfic treatment cells were treated with 10 nM okadaic acid (OA) 16 hours prior to collection. The medium was removed 72 hours post-transfection followed by cell fixation using 4% paraformaldehyde solution (BD Biosciences) at 4^*°*^C for 20 minutes. After fixation, the samples were immersed in PBS and stored at 4^*°*^C until confocal microscopy analysis.

### HA-Tag Immunoprecipitation

HEK293T cells were co-transfected with constructs HA-Rab8a-CA (Q67L, Addgene, Cat # 101050), 2xMyc-LRRK2-Y1699C (Addgene, Cat # 25364), and PepRePs with Opti-MEM and Lipofectamine 2000 for 48 hours prior to harvest. After treatment, cells were lysed in 500 *µ*L lysis buffer (150 mM NaCl, 1 mM EDTA, 25 mM Tris-HCl, 1% NP-40), collecting lysates by pipetting. Each lysate was passed through a 23-gauge needle seven times then centrifuged at 15,000 rpm for 20 minutes. Protein concentration was determined by Pierce BCA Protein Assay Kit (ThermoFisher, Cat # 23227). HA-tag beads were pre-washed with lysis buffer before adding 1 mg of protein lysate, where the total volume was adjusted to 500 *µ*L with lysis buffer. Samples were incubated for 16 hours at 4^*°*^C with rotation; subsequently, beads were collected on a magnetic stand and were subjected to six wash steps with 500 *µ*L wash buffer (75 mM NaCl, 0.5 mM EDTA, 12.5 mM Tris-HCl, 0.5% NP-40), inverting the tube several times and incubating on ice for 5 minutes per wash step. Bound proteins were eluted in 20 *µ*L 1X SDS sample buffer (60 mM Tris-HCl [pH 6.8], 10% glycerol, 2% SDS, 0.1% NaN_3_, 1% 2-mercaptoethanol, 0.01% bromophenol blue) and boiled at 100^*°*^C for 5 minutes prior to separation on an 8% SDS-PAGE gel.

### Immunoprecipitation-Mass Spectrometry (IP-MS)

HEK293T cells were transfected with HA-Rab8A, the indicated PepRePs constructs, and Myc-LRRK2 using Opti-MEM and Lipofectamine 2000. For pharmacological inhibition experiments, cells were treated with MLi-2 prior to harvest. Cells were collected and lysed in lysis buffer containing 150 mM NaCl, 1 mM EDTA, 25 mM Tris-HCl, 1% (v/v) NP40, phosphatase inhibitor (PhosStop, Roche), and protease inhibitor (cOmplete, Roche). Lysates were clarified by centrifugation at 15000 rpm for 20 minutes at 4^*°*^C, and the resulting supernatants were collected for immunoprecipitation. HA-Rab8A-containing complexes were immunoprecipitated using anti-HA affinity beads and subsequently washed to remove nonspecific proteins. Immunoprecipitated samples were processed for liquid chromatography-tandem mass spectrometry (LC-MS/MS) analysis. Protein abundance was quantified across experimental conditions and normalized to Rab8A bait abundance. Differentially associated proteins were identified by comparison of MLi-2-treated and PepReP-expressing samples to the corresponding control conditions. Concordance of treatment effects between MLi-2-mediated LRRK2 inhibition and Rab8A_SnP_2-PP1c-mediated Rab8A dephosphorylation was assessed by comparing bait-normalized protein abundance across conditions.

### Immunoblotting

On the day of harvest, cells were detached by adding 0.5% trypsin-EDTA then washed twice with ice-cold 1X DPBS. Cells were then lysed using a 1:100 dilution of Halt™ Protease and Phosphatase Inhibitor Cocktail (100X, ThermoFisher, Cat # 78440) in Pierce RIPA buffer (ThermoFisher, Cat # 89900). Supernatants containing whole-cell protein samples were quantified using the Pierce BCA Protein Assay Kit (ThermoFisher, Cat # 23227); twenty micrograms per sample was mixed with 4X Bolt™ LDS Sample Buffer (ThermoFisher, Cat # NP0007) and 5% 2-mercaptoethanol (Millipore Sigma, Cat # M3148) prior to immunoblotting, which was performed according to standard protocols. Proteins were probed with mouse anti-*β*-actin antibody (Cell Signaling Technology, Cat # 3700), mouse anti-GAPDH (Santa Cruz, Cat # sc-47724), rabbit anti-vinculin (ThermoFisher, Cat # 700062), rabbit anti-tubulin antibody (Cell Signaling Technology, Cat # 2144S), mouse anti-tau antibody (Santa Cruz, Cat # sc-21796), mouse anti-pT231 tau antibody (ThermoFisher, Cat # MN1040), rabbit anti-Rab8A antibody (Cell Signaling Technology, Cat # 6975), or rabbit anti-pT72 Rab8A (Abcam, Cat # MJF-R20) overnight at 4^*°*^C. The blots were washed and probed with secondary antibodies for 1-2 hours at room temperature before imaging. Densitometry analysis of protein bands in immunoblots was performed using FIJI software as described here: (https://imagej.nih.gov/ij/docs/examples/dot-blot/). Intensity data for the PepRePs bands was first normalized to band intensity of *β*-actin in each lane, then to the average band intensity of the whole protein, then to the non-targeting control cases across replicates.

### High-Throughput Immunoblotting

For higher-throughput quantification, we measured STAT3 pS727 and PTEN pS380 using the Lumit® Immunoassay Cellular System (Promega, Cat # W1201) in 96-well format, plating 20,000 cells per well: MDA-MB-231 cells for STAT3, and A549 cells for PTEN. For each target, we paired a mouse antibody against the total protein with a rabbit antibody against the phosphorylated form: STAT3 (Cell Signaling Technology, Cat # 9139S) with phospho-STAT3 Ser727 (Cell Signaling Technology, Cat # 9134S) and PTEN (Cell Signaling Technology, Cat # 9188S) with phospho-PTEN Ser380 (CST, Cat # 9549S), each used at the dilution recommended in the corresponding Promega Application Note. We combined the cell lysate, antibody mix, and Lumit Detection Substrate at a 50 *µ*L:50 *µ*L:25 *µ*L ratio per well following the manufacturer’s protocol, then read luminescence on a Promega GloMax® plate reader. Three replicates were performed per condition, with luminescence normalized to a non-targeting loading control.

### MTS Proliferation Assay

We seeded A549 cells (for PTEN constructs) or MDA-MB-231 cells (for STAT3 constructs) into 96-well plates at 3,000 cells per well in complete growth medium (DMEM or RPMI-1640, 10% FBS, 1% penicillin-streptomycin) and allowed cells to adhere for 24 h at 37^*°*^C before transfection or treatment. At each time point (0, 6, 12, 24, and 48 h post-treatment), we added 20 *µ*L of CellTiter 96® AQueous One Solution Reagent (Promega, Cat # G3580) directly to each well containing 100 *µ*L of medium, without removing or replacing existing medium, and returned plates to the incubator for 1–2 h. We recorded absorbance at 490 nm on a plate reader, using wells containing medium and MTS reagent but no cells as a blank, and subtracted the mean blank value from all readings before analysis. Each condition was assayed in triplicate wells per time point.

### Statistical Analysis and Reproducibility

Sample sizes were not pre-determined by power analysis; three independent biological replicates were used per condition, consistent with standards of the field. All data was reported as mean values, with error bars representing standard deviation. Statistical comparisons used a two-tailed Student’s test, ordinary one-way ANOVA with Dunnett’s multiple comparisons tests, and two-way ANOVA with Dunnett’s multiple comparisons tests using GraphPad Prism 11 software: ^*^, p *≤*0.05; ^**^, p *≤*0.01; ^***^, p *≤*0.001; ^****^, p *≤*0.0001. P-values shown above bars in densitometry analyses and other experiments compare samples to the control. No data were excluded from the analyses. Experiments were not randomized, and investigators were not blinded to allocation or outcome assessment.

## 5 Author Contributions

L.H. designed the PepRep architecture and constructs and conducted PTEN and STAT3 experiments, with input from T.C.D and M.P.D. T.C.H. Y.S.T. performed all tau and Rab8a experiments, with supervision from P.H.C. S.V. and T.C. performed computational design of all guide peptides. P.C. conceived, designed, and directed the study, and reviewed and finalized the manuscript.

## 6 Data Availability

All data used in this paper are present in the paper, supplementary tables, and supplementary figures. Raw and processed data are provided as Source Data in our public Zenodo repository: https://doi.org/10.5281/zenodo.22883270. The PepRePs cloning vector will be deposited to Addgene upon publication.

## 7 Acknowledgments

This research was supported by NIH grant R35GM155282. We also thank UbiquiTx, Inc. for access to the core SaLT&PepPr codebase.

## Supplementary Information

**Figure S1:**
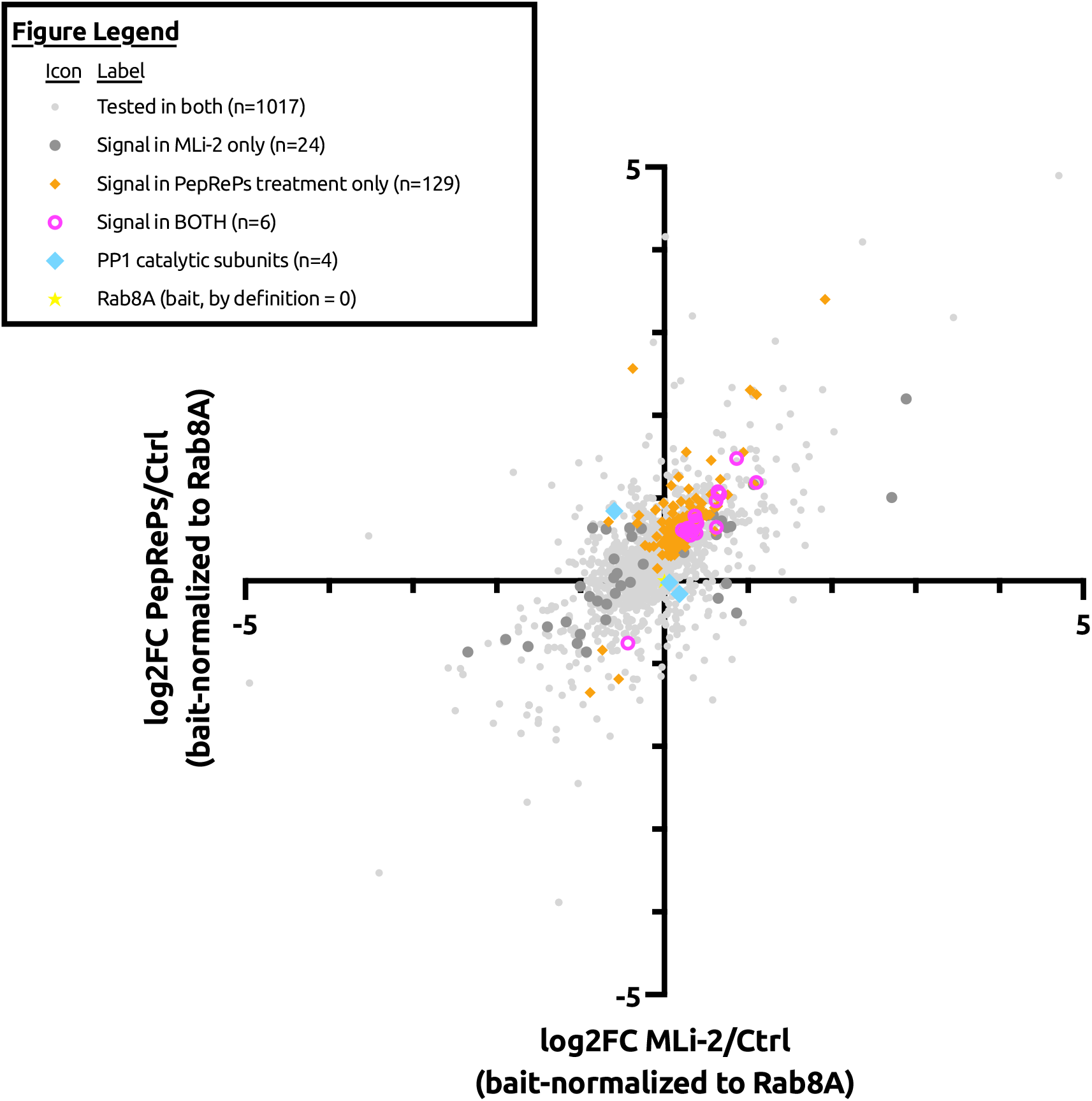
Correlation between PepRePs-mediated and pharmacological perturbation of the Rab8A interactome. Pearson and Spearman correlation analysis of changes in Rab8A-associated proteins following Rab8A_SnP_2-PP1c expression or MLi-2 treatment. Each point represents a protein quantified by quantitative proteomic analysis following HA-tag enrichment of Rab8A-associated complexes. Correlation was calculated using the treatment-associated change in protein abundance for each perturbation. Pearson *r* = 0.67 and Spearman *ρ* = 0.64.

**Table S1:**
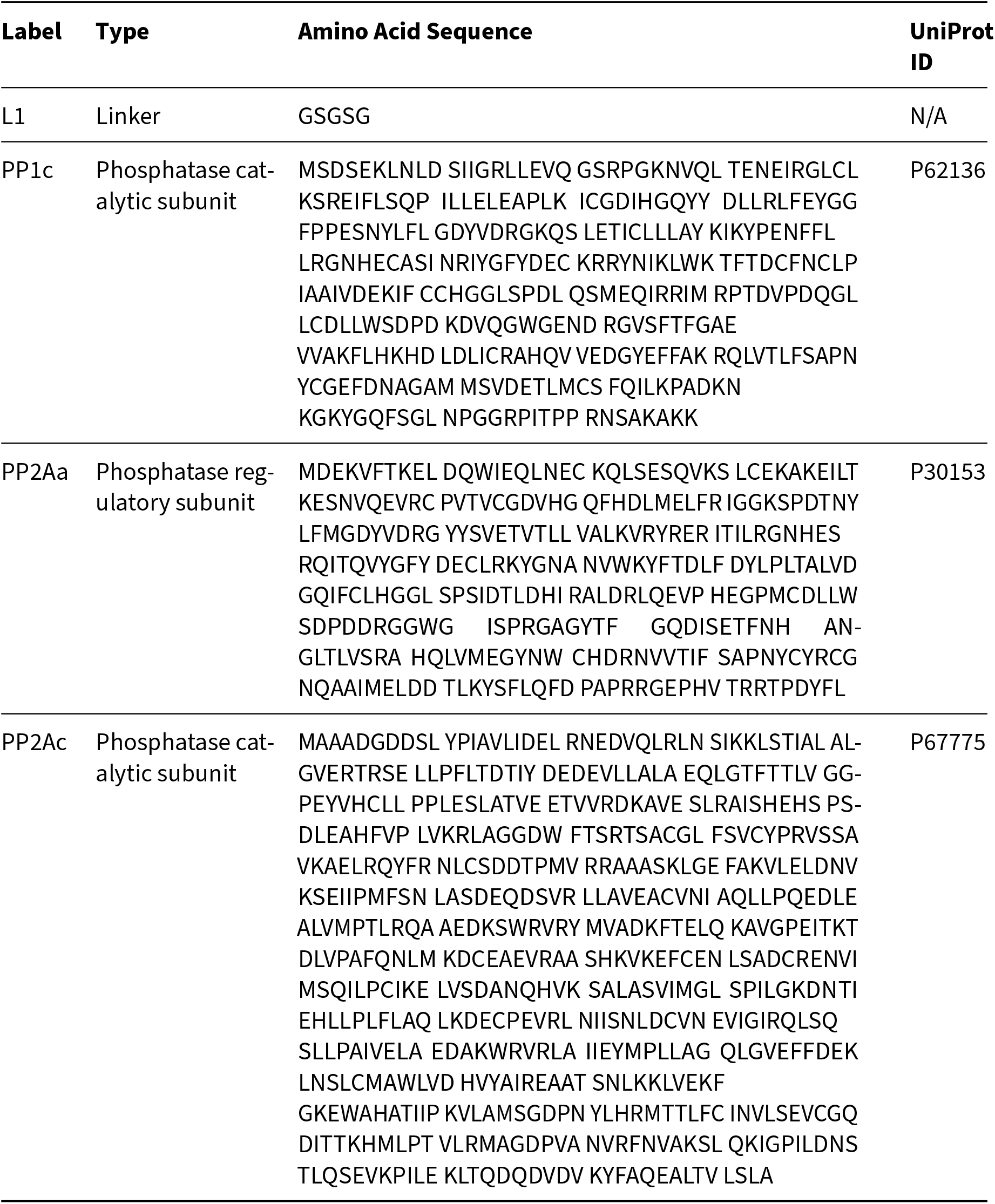
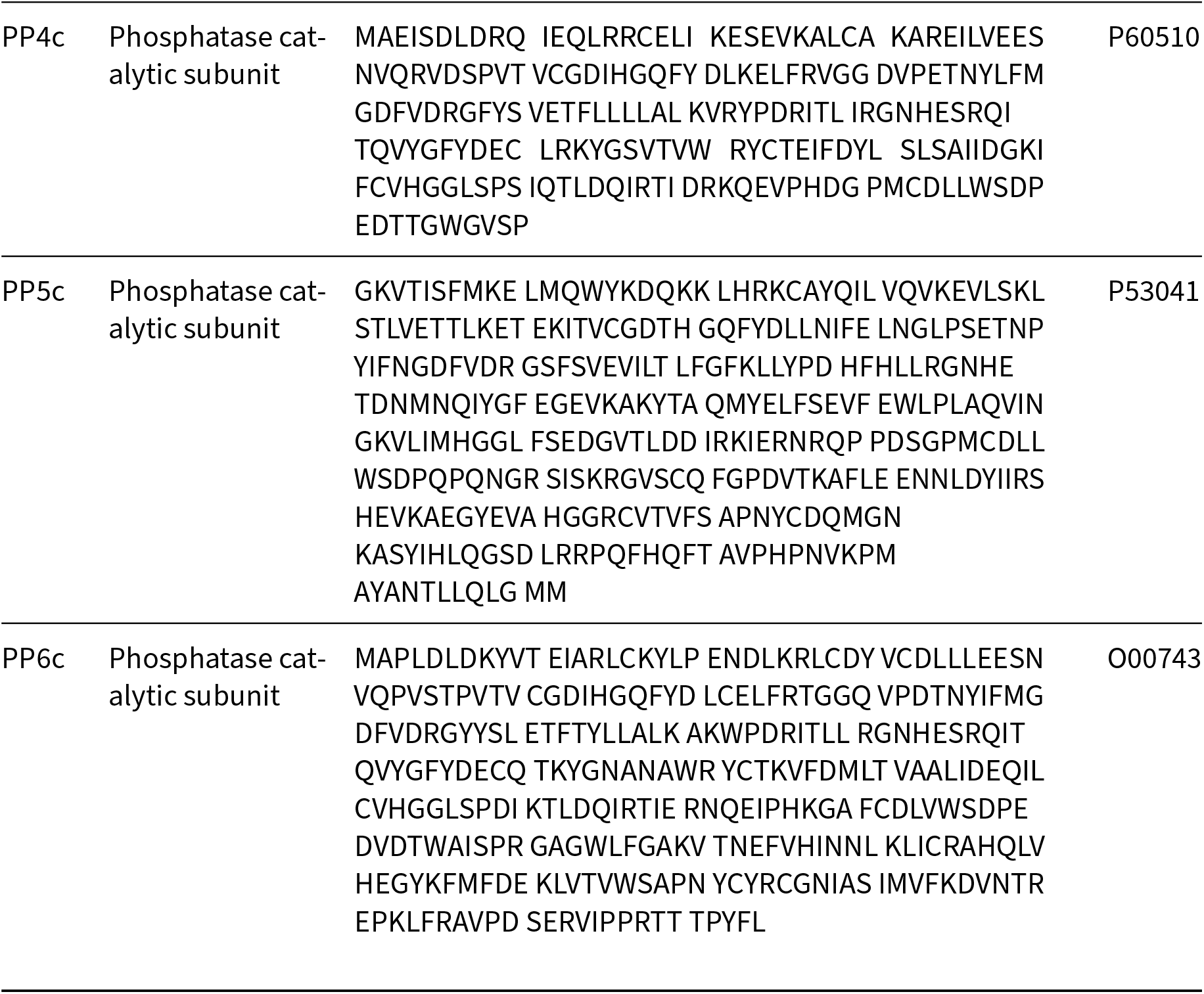
Phosphatase subunit and linker sequences used in this study.

**Table S2:**
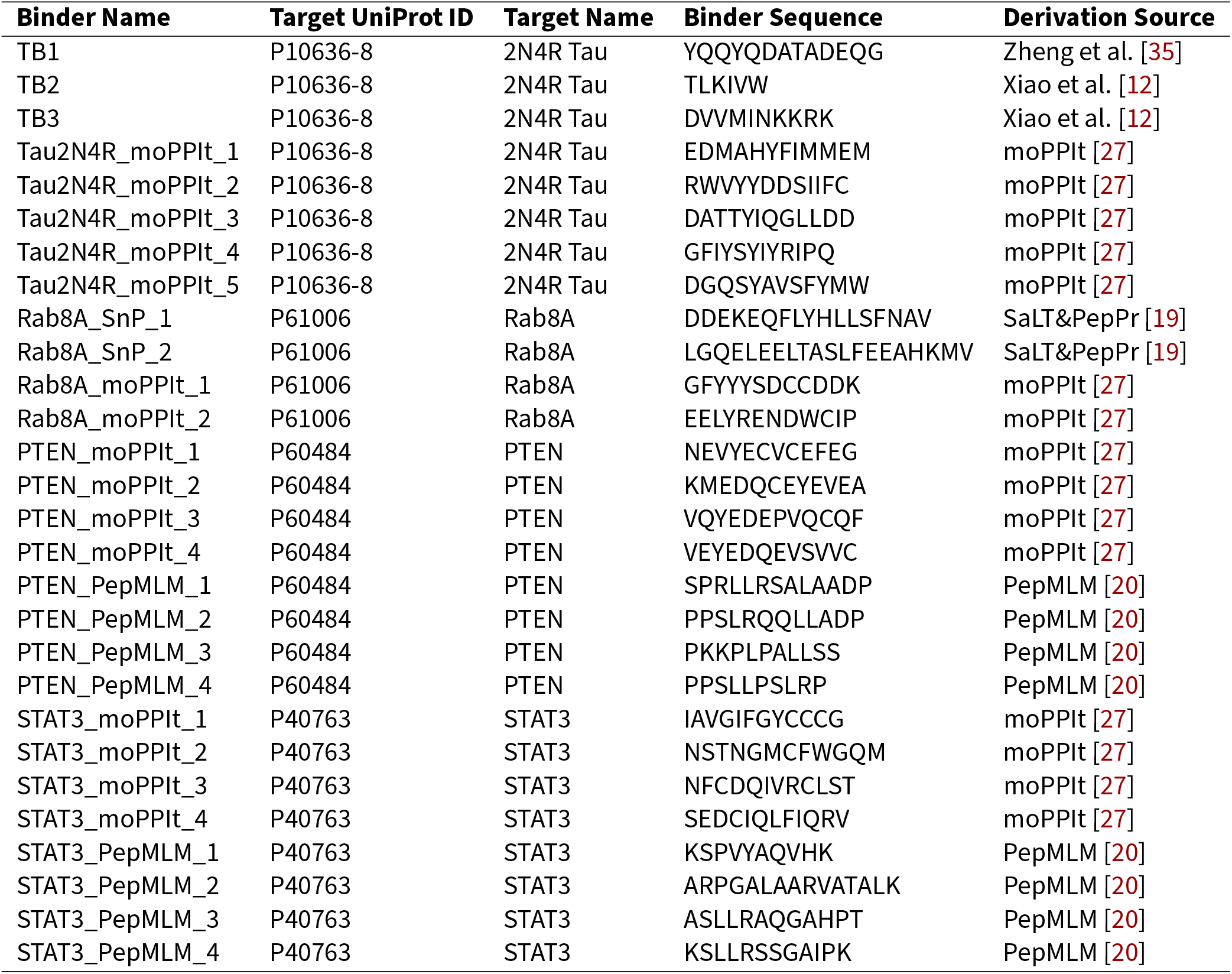
Peptide binders used in this study.

